# Securing the exchange of synthetic genetic constructs using digital signatures

**DOI:** 10.1101/750927

**Authors:** Jenna E Gallegos, Diptendu M. Kar, Indrakshi Ray, Indrajit Ray, Jean Peccoud

## Abstract

Synthetic biology relies on an ever-growing supply chain of synthetic genetic material. Technologies to secure the exchange of this material are still in their infancy. Solutions proposed thus far have focused on watermarks, a dated security approach that can be used to claim authorship, but is subject to counterfeit, and does not provide any information about the integrity of the genetic material itself. We describe how data encryption and digital signature algorithms can be used to ensure the integrity and authenticity of synthetic genetic constructs. Using a pilot software that generates digital signatures and other encrypted data for plasmids, we demonstrate that we can predictably extract information about the author, the identity, and the integrity of plasmid sequences from sequencing data alone without a reference sequence, all without compromising the function of the plasmids. We discuss how this technology can be improved, applied, and expanded to support the new bioeconomy.

## Introduction

Synthetic biologists use software tools to generate DNA sequences encoding complex functions(*1-4*). In this context, watermarks (*5-8*) have been inserted in synthetic DNA to assert authorship claims. DNA watermarking demonstrates the need to assert authorship, but watermarks are insufficient security solutions. We propose the application of digital signatures to DNA in order to support more trustworthy transactions in the bioeconomy supply chain (*9, 10*).

A digital signature is a mathematical technique used to validate digital messages, documents, web pages, and software. The signature of a document relies on a hash function that maps the entire document to a small string of a set length. The resulting hash value is then encrypted using the author’s private key. The person receiving the document can decrypt the signature using the author’s public key. Therefore, unlike watermarks, digital signatures are derived from and inextricably linked to both the content being signed and the author. Digital signatures ensure authenticity by enabling the recipient to confirm that the content was generated by an authentic source. Digital signatures also ensure integrity, because the signature is invalidated if the signed content is changed in any way.

As electronic documents are signed by embedding digital signatures, it is conceptually possible to digitally sign genetic constructs by inserting DNA sequences encoding the output of a digital signature algorithm. Such an approach would provide a greater level of security than watermarks, but is impractical for three reasons:

1. Validation would require some knowledge of the author’s identity.
2. This approach would not allow for unique identification of the genetic construct itself.
3. The signature validation algorithm would invalidate any sequence that does not perfectly match that which was signed. This stringent validation may be excessive considering that some mutations may not compromise the construct’s behavior.

## Results

### Signature Generation and Verification

To address these challenges, we developed a system for applying digital signatures to DNA that is robust to minor mutations and establishes authorship and sequence identification without prior knowledge using the following innovations:

1. We used the author’s ORCID as a public key and encoded it in the DNA sequences. Other public identifier’s like the DUNS number of companies could also be used.
2. We encoded in the sequence a six-digit user-generated number, which serves as a plasmid identifier (Plasmid ID). The Plasmid ID could correspond to a laboratory information management system (LIMS) record, a serial number for a licensing agreement or a catalog number.
3. We introduced an Error Correction Code (ECC) that makes it possible to identify a limited number of mutationsin the event of an invalidated signature. The ECC relies on algorithms like those that use redundancy in DVD’s to reconstruct the original data. The 32 base pair ECC used in the subsequently described experiments detects up to two single nucleotide polymorphisms (SNPs).

We developed a pilot software that takes a GenBank (.gb) file as input along with the Author’s ORCID and a Plasmid ID. The application generates a new GenBank file that includes the ORCID, Plasmid ID, ECC, and the digital signature at a user-defined location. The algorithms used for the software are described in detail in a previous publication(*11*). This “signed” plasmid can then be ordered from a DNA synthesis company or assembled from individual DNA fragments.

Using the same software, plasmid users can recover the ORCID and Plasmid ID from assembled sequencing reads in the form of a FASTA file. To identify the signature cassette within the sequence, the software uses 10bp sequence delimiters. The software identifies the signature cassette, extracts the author and plasmid identities, and determines whether or not the signature is valid by regenerating the original sequence from the decrypted signature. If the signature is invalid, the ECC is invoked. If the number of errors is below the threshold set by the ECC, the software reports the changes. If the number of errors exceeds the threshold, the software reports that there are too many errors.

This system thus allows the recipient of a plasmid to verify that the sample was shared by an authentic sender and that the original sequence is unchanged, or changed only in trivial ways, all without a reference sequence (Figure 1).

**Figure 1.**
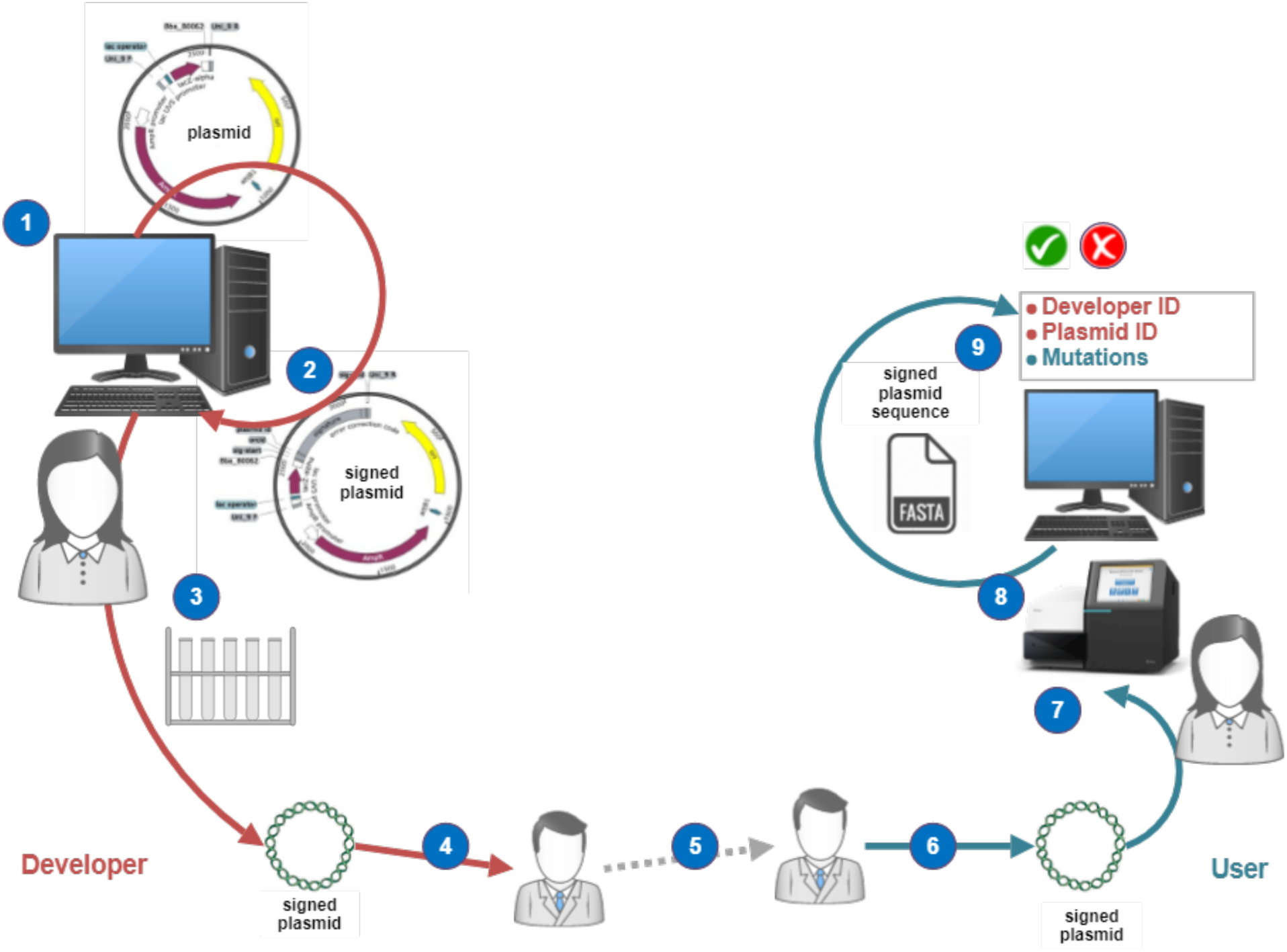
Secured exchange of signed plasmids. (1) Developer designs plasmid; (2) Using software, developer signs the plasmid with private key. Signature is inserted in the plasmid sequence along with the developer’s ORCID, plasmid ID, and ECC. (3) Developer synthesizes signed plasmid from its computer-generated sequence. (4-5) Signed plasmid circulates through a series of undocumented transactions. (6) Plasmid user receives the signed plasmid. (7) Plasmid user sequences plasmid. (8) Sequencing reads are assembled to generate the complete unannotated sequence of the signed plasmid. (9) Software extracts the ORCID, plasmid ID, and detects mutations. Using the ORCID as public key, software validates the digital signature, ensuring the integrity and authenticity of the signed plasmid.

## Experimental Validation

To validate our ability to verify digital signatures from sequencing data and ensure that digital signatures do not interfere with the function of plasmids, a series of plasmids were designed. These plasmids are described in Figure 2 and include:

**Figure 2.**
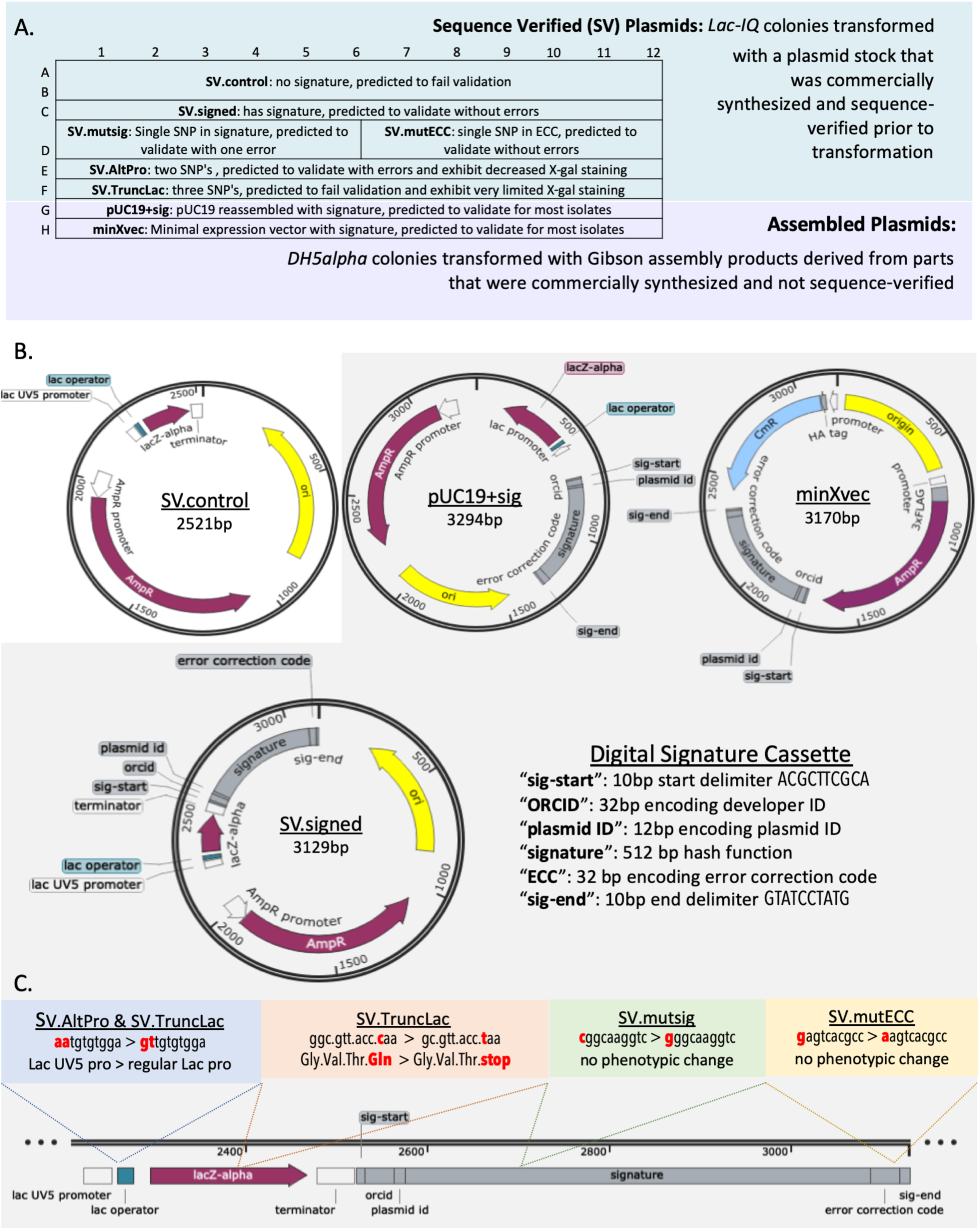
Description of plasmids in this study. **A)** Depicts how transformants of each plasmid are arrayed on a 96-well plate and describes expected result. Includes 24 transformants for SV.control; 12 for SV.signed, SV.AltPro, and SV.TruncLac; 6 for SV.mutsig and SV.mutECC; and, 12 isolates from Gibson assemblies for pUC19+sig and minXvec. **B)** Maps for four of the eight plasmids and a breakdown of digital signature cassette components. **C)** Schematic showing how each of the remaining four plasmids differs from SV.signed.

1. A commercial plasmid (pUC19+sig)
2. A minimal expression vector (minXvec)
3. A family of six sequence verified clones with and without signatures with varying mutations (SV.control, SV.signed, SV.mutsig, SV.mutECC, SV.AltPro, and SV.TruncLac).

Plasmids pUC19+sig and minXvec were constructed by Gibson assembly(*12*) of four DNA fragments. Because the fragments were not sequence-verified, some random mutations were expected.

The sequence verified plasmids include a *lacZ* expression construct. These constructs were ordered from Twist Biosciences as sequence verified clones within one of the vendor’s predefined backbones. SV.control and SV.signed are identical except that the latter contains a signature cassette.

SV.mutsig and SV.mutECC each have a SNP in the signature or ECC, respectively. SV.AltPro has two SNPs that convert the stronger *lac*UV5 promoter (*13*) to the weaker wildtype *lac* promoter. SV.TruncLac is the same as SV.AltPro except for an additional SNP that introduces a premature stop codon after the 24^th^ amino acid of the *lacZ* gene.

DNA was extracted from 6-24 transformants for each plasmid for a total of 96 samples and sequenced. Using the assembled FASTA files, signature validation was performed. The results for each plasmid were confirmed via manual reference alignment and are represented in Figure 3a.

**Figure 3.**
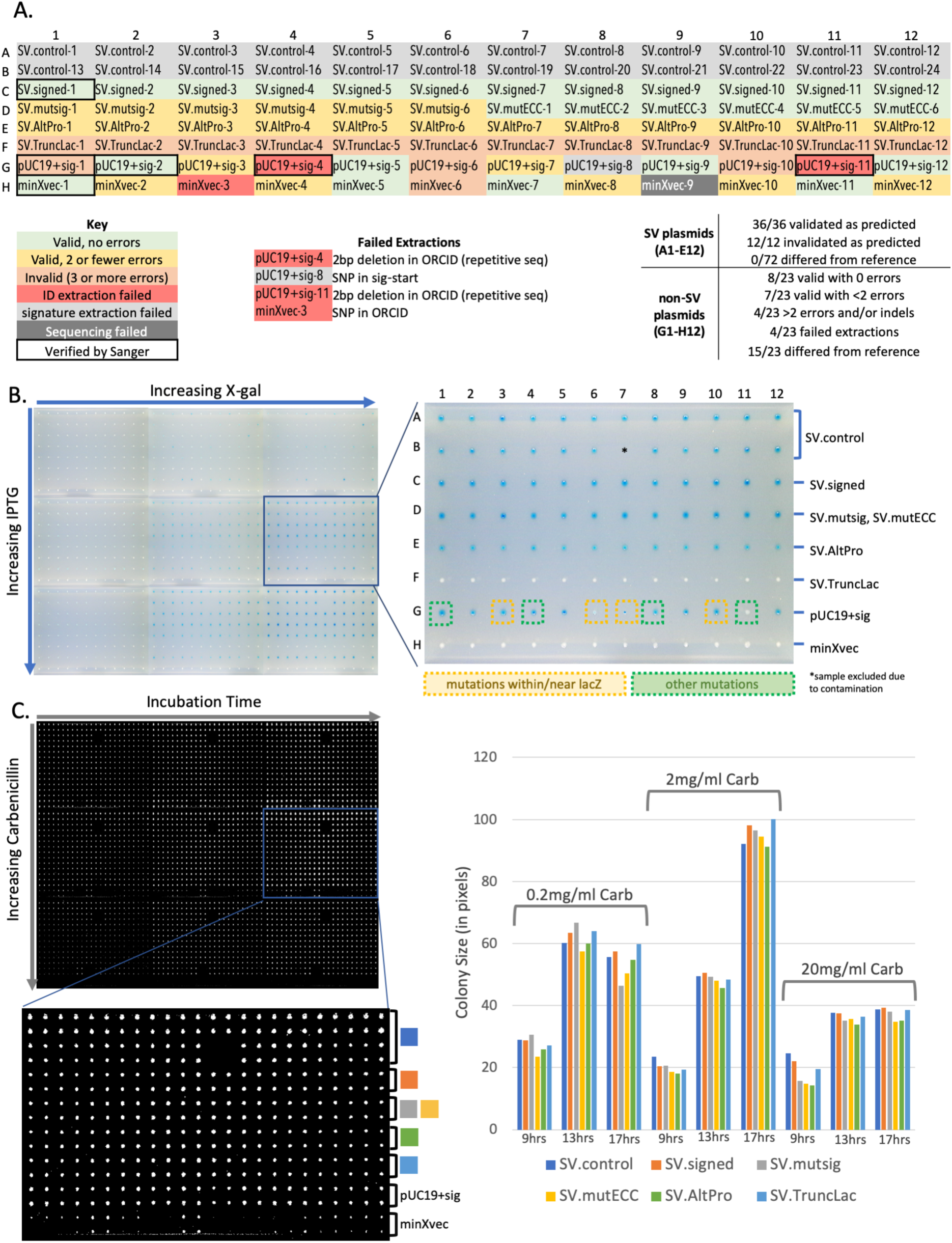
Signature validation and phenotyping outcomes. **A)** Signature validation for each of the 96 samples. **B)** Comparison of lacZ expression in each of the 96 samples across variable inducer (IPTG) and substrate (X-gal) concentrations after 10 hours of incubation. **C)** Comparison of colony growth for each of the 96 samples pinned in quadruplicate on variable concentrations of carbenicillin up to 20mg/ml.

All 72 of the transformants from the sequence verified family of plasmids passed/failed validation as expected. 15/23 of the isolates transformed with the Gibson-assembled plasmids (pUC19+sig and MinXvec) contained mutations. Seven of these had two or fewer SNPs, which were captured by the validation software. Four of these with more than two SNPs and/or insertions/deletions (INDELs) failed validation. The remaining four had mutations in the signature cassette itself which resulted in failed signature or ID extraction.

To determine whether adding 608bp of extraneous sequence negatively impacted the function of the plasmids, we replica plated the 96 transformants onto media with varying concentrations of antibiotic, the *lacZ* substrate (X-gal), or lac operator inducer (IPTG).

The sequence verified family of plasmids were all expected to exhibit comparable, inducible levels of *lacZ* expression, except for SV.AltPro, which has a weaker promoter(*13*) and SV.TruncLac, which has a weaker promoter and a premature stop codon(*14*).

The addition of the digital signature cassette did not noticeably impact antibiotic resistance, *lacZ* expression, or induction (Figure 3). Visible sources of variability in expression included: promoter conversion and protein truncation in SV.AltPro and SV.TruncLac, random mutations in the non-sequence verified parts used to assemble pUC19+sig and minXvec, and random *lacZ* induction in the absence of IPTG, which occurred in some colonies regardless of the presence/absence of a signature.

Otherwise, individual transformants containing the same plasmid performed comparably within the sequence-verified plasmid family, indicating that the signature cassette is unlikely to impact plasmid function, stability or retention in the timescale of the tests performed.

The only transformant that did not behave as expected is pUC19+sig-11 (Figure 3b). It is unclear why *lacZ* expression was silenced in this case, but the plasmid is identical to pUC19+sig-4 except that pUC19+sig-4 contains an additional SNP in the signature, so the presence of the signature is likely irrelevant.

## Discussion

In this manuscript, we’ve demonstrated the feasibility of using digital signatures to encrypt and secure DNA in living organisms.

Our system is robust. We validated/invalidated 72 sequence-verified clones with 100% predictability, indicating that neither DNA synthesis, nor sequencing, nor sequence assembly will present significant technical barriers to employing digital signature-based verification schemes for plasmid sequences.

Our system is sensitive. We identified 30 isolates with known mutations and 14 with unknown mutations including single SNPs and small INDELS without the use of a reference sequence. Out of 96 samples, our system performed as intended in all but five instances (discussed below).

Finally, introducing a 608bp signature cassette into a ∼2.5kb plasmid did not have obvious detrimental effects. Nonetheless, we are working to optimize compression algorithms to further decrease the length of the signature (*15*).

Of the five samples for which we did not achieve the expected result, four had mutations within the signature cassette. In one case (pUC19+sig-8), a mutation in the start delimiter caused signature extraction to fail. We have already developed a solution to this problem, in which the software searches not only for the exact start sequence, but also for related sequences(*15*).

In three cases (pUC19+sig-4, pUC19+sig-11, and minXvec-3), a mutation in the ORCID or plasmid ID caused the pilot software to return incorrect information regarding the identify/origin of the sample. Cases where one of the ID strings contains a SNP (ex. minXvec-3) can be amended by revising the pilot software so that it returns an updated ORCID and plasmid ID following error correction. INDELs are much more challenging computationally.

When performing validation, our pilot software aligns the original sequence (recovered from the hash) with the sequence submitted for validation, using the start delimiter as an origin. Hence SNPs can be identified by noting any position where the original and current sequences do not match. In future iterations, we could incorporate more complex sequence alignment algorithms to identify INDELs during error correction.

One additional sample for which we did not achieve the expected result (minXvec-9) simply was not assembled due to the low quality of the sequencing reads. High quality assemblies for 95/96 plasmid samples is a better-than-expected result, especially considering that most of the plasmids submitted (the entire sequence verified family) had the capacity to form a substantial hairpin. We found that we could sequence each of our ∼3kb test plasmids for $32 by Sanger sequencing (not including primers) or $35 by Illumina (including library prep and sequence assembly for a minimum of 96 samples). Continually decreasing sequencing costs and affordable, primer-agnostic technologies such as Oxford Nanopore will likely facilitate the adoption of our approach.

## Methods

### Plasmid Construction and Sequencing

The sequence-verified family of plasmids was designed using GenoCAD software (https://design.genofab.com/). The vector backbone and inserts were synthesized and sequence-verified by Twist Biosciences (www.twistbiosciences.com).

pUC19+sig and minXvec were each constructed by Gibson assembly(*12*) of four DNA fragments synthesized by either Integrated DNA Technologies (www.idtdna.com) or Twist Biosciences (www.twistbiosciences.com) based on price and technical limitations. Parts for Gibson assembly of pUC19+sig and minXvec were designed with 40bp overlaps. Plasmids were assembled using a Gibson Assembly^®^ HiFi 1 Step Kit (Synthetic Genomics, Inc., La Jolla, CA), according to the manufacturer’s instructions.

DNA was extracted from each of the 96 isolates using a ZR Plasmid Miniprep – Classic Kit (Zymo Research, Irvine, CA). After quality control analysis on an Agilent TapeStation, ∼15-150ng of each isolate was submitted to SeqWell in a 96-well plate. The plasmids were sequenced at 90x coverage. Assembled FASTA files provided by SeqWell from their proprietary plasmid sequence assembly platform were used for validation.

### Phenotyping

NEB^®^ 5-alpha F’Iq Competent E. coli (SV plasmids) or NEB^®^ 5-alpha Competent E. coli (pUC19+sig and minXvec) cells from which DNA was extracted for sequencing were pinned from frozen stocks onto LB+100ug/ml Carbenicillin plates using a Rotor HAD robot (Singer Instruments, Somerset, UK). Colonies were grown at 37 degrees to saturation and stored in the fridge.

“Master plates” were generated by pinning from the plates in the fridge onto fresh LB+100ug/ml carbenicillin plates in either 96 (for variable X-gal/IPTG) or 384 (for variable carbenicillin) array. After ∼12 hours at 37 degrees, these plates were each pinned onto the phenotyping plates. A different master plate was used for every 3-6 phenotyping plates.

Phenotyping plates were all standard LB with variable concentrations of carbenicillin (carb), 5-bromo-4-chloro-3-indolyl-β-D-galactopyranoside (X-gal), or Isopropyl β-D-1-thiogalactopyranoside (IPTG). Variable carb plates contained 100ug/ml, 200ug/ml, 1mg/ml, 2mg/ml, 1-mg/ml, or 20mg/ml antibiotic. Variable IPTG/X-gal plates contained 100ug/ml carb, 0ul/ml, 0.1ul/ml, or 1ul/ml 100mM IPTG, and 1ul/ml, 5ul/ml, or 10ul/ml 50mM X-gal.

Plates were imaged over two-hour time courses using a Phenobooth (Singer Instruments, Somerset, UK) imaging platform. Plates which best demonstrated any differences between the colonies were selected for figures. To generate the graph in Figure 3C, Phenobooth imaging software was used to process images and export colony size data. Quadruplicate technical replicates of each isolate were averaged, and then those values were averaged across each of the biological replicates (between six and 23) for a given time point on a given plate.

The technology we’ve described is a first step towards management of intellectual property (IP) embedded in genetic constructs. This approach does not provide as much security as a software key, because the biological function of the sequences is not dependent on the signature/key. However, it facilitates several facets of IP protection. In addition to sequence and author verification, this technology allows for the serialization of genetic constructs, making it possible to uniquely identify genetic material distributed to different users and the licensing agreements they are associated with. The technology can also be used for attribution in cases of leaked genetic material with IP or security implications, such as infectious agents(*8, 16*) or controlled substances like biologically-derived opioids(*17*).

We anticipate that authentication of DNA sequences and their authors will become increasingly relevant as DNA is used for information sharing and storage in the future(*18, 19*).

## Methods

### Plasmid Construction and Sequencing

The sequence-verified family of plasmids was designed using GenoCAD software (https://design.genofab.com/). The vector backbone and inserts were synthesized and sequence-verified by Twist Biosciences (www.twistbiosciences.com).

pUC19+sig and minXvec were each constructed by Gibson assembly(*12*) of four DNA fragments synthesized by either Integrated DNA Technologies (www.idtdna.com) or Twist Biosciences (www.twistbiosciences.com) based on price and technical limitations. Parts for Gibson assembly of pUC19+sig and minXvec were designed with 40bp overlaps. Plasmids were assembled using a Gibson Assembly^®^ HiFi 1 Step Kit (Synthetic Genomics, Inc., La Jolla, CA), according to the manufacturer’s instructions.

DNA was extracted from each of the 96 isolates using a ZR Plasmid Miniprep – Classic Kit (Zymo Research, Irvine, CA). After quality control analysis on an Agilent TapeStation, ∼15-150ng of each isolate was submitted to SeqWell in a 96-well plate. The plasmids were sequenced at 90x coverage. Assembled FASTA files provided by SeqWell from their proprietary plasmid sequence assembly platform were used for validation.

### Phenotyping

NEB^®^ 5-alpha F’Iq Competent E. coli (SV plasmids) or NEB^®^ 5-alpha Competent E. coli (pUC19+sig and minXvec) cells from which DNA was extracted for sequencing were pinned from frozen stocks onto LB+100ug/ml Carbenicillin plates using a Rotor HAD robot (Singer Instruments, Somerset, UK). Colonies were grown at 37 degrees to saturation and stored in the fridge.

“Master plates” were generated by pinning from the plates in the fridge onto fresh LB+100ug/ml carbenicillin plates in either 96 (for variable X-gal/IPTG) or 384 (for variable carbenicillin) array. After ∼12 hours at 37 degrees, these plates were each pinned onto the phenotyping plates. A different master plate was used for every 3-6 phenotyping plates.

Phenotyping plates were all standard LB with variable concentrations of carbenicillin (carb), 5-bromo-4-chloro-3-indolyl-β-D-galactopyranoside (X-gal), or Isopropyl β-D-1-thiogalactopyranoside (IPTG). Variable carb plates contained 100ug/ml, 200ug/ml, 1mg/ml, 2mg/ml, 1-mg/ml, or 20mg/ml antibiotic. Variable IPTG/X-gal plates contained 100ug/ml carb, 0ul/ml, 0.1ul/ml, or 1ul/ml 100mM IPTG, and 1ul/ml, 5ul/ml, or 10ul/ml 50mM X-gal.

Plates were imaged over two-hour time courses using a Phenobooth (Singer Instruments, Somerset, UK) imaging platform. Plates which best demonstrated any differences between the colonies were selected for figures. To generate the graph in Figure 3C, Phenobooth imaging software was used to process images and export colony size data. Quadruplicate technical replicates of each isolate were averaged, and then those values were averaged across each of the biological replicates (between six and 23) for a given time point on a given plate.

## Acknowledgements

This work was supported by NSF award #1934573 “EAGER: Development of a tool-chain to write and read self-documenting plasmids”, NSF award #1832320 “EAGER: Modeling DNA Manufacturing Processes Using Extensible Attribute Grammars”, and by Colorado State University’s Office of the Vice President for Research Catalyst for Innovative Partnerships Program. The content is solely the responsibility of the authors and does not necessarily represent the official views of the Office of the Vice President for Research. The work of Indrajit Ray was performed while serving as Program Director at the U.S. National Science Foundation (NSF) and supported by the foundation’s Independent Research and Development program for staff. Research findings presented here, and opinions expressed are solely that of the authors, and in no way reflect the opinion of the NSF or other federal agencies.

## References

1. A. A. Nielsen et al., Genetic circuit design automation. Science 352, aac7341 (2016).

2. H. M. Salis, E. A. Mirsky, C. A. Voigt, Automated design of synthetic ribosome binding sites to control protein expression. Nature Biotechnology 27, 946 (2009).

3. A. Villalobos, J. E. Ness, C. Gustafsson, J. Minshull, S. Govindarajan, Gene Designer: a synthetic biology tool for constructing artificial DNA segments. BMC Bioinformatics 7, 285 (2006).

4. P.-S. Huang, S. E. Boyken, D. Baker, The coming of age of de novo protein design. Nature 537, 320 (2016).

5. M. Liss et al., Embedding Permanent Watermarks in Synthetic Genes. Plos One 7, (2012).

6. D. Heider, A. Barnekow, DNA-based watermarks using the DNA-Crypt algorithm. BMC Bioinformatics 8, 176 (2007).

7. G. Thouand, R. Marks, Bioluminescence: Fundamentals and Applications in Biotechnology - Volume 3 Preface. Adv Biochem Eng Biot 154, V-V (2016).

8. D. C. Jupiter, T. A. Ficht, J. Samuel, Q.-M. Qin, P. De Figueiredo, DNA watermarking of infectious agents: progress and prospects. PLoS pathogens 6, e1000950 (2010).

9. J. Peccoud, J. E. Gallegos, R. Murch, W. G. Buchholz, S. Raman, Cyberbiosecurity: From Naive Trust to Risk Awareness. Trends Biotechnol 36, 4–7 (2018).

10. R. S. Murch, W. K. So, W. G. Buchholz, S. Raman, J. Peccoud, Cyberbiosecurity: An Emerging New Discipline to Help Safeguard the Bioeconomy. Front Bioeng Biotechnol 6, 39 (2018).

11. D. M. Kar, I. Ray, J. Gallegos, J. Peccoud, paper presented at the Proceedings of the New Security Paradigms Workshop, Windsor, United Kingdom, 2018.

12. D. G. Gibson et al., Enzymatic assembly of DNA molecules up to several hundred kilobases. Nat Methods 6, 343–U341 (2009).

13. R. J. Noel, W. S. Reznikoff, Structural studies of lacUV5-RNA polymerase interactions in vitro - Ethylation interference and missing nucleoside analysis. Journal of Biological Chemistry 275, 7708–7712 (2000).

14. K. Nishiyama, N. Ichihashi, Y. Kazuta, T. Yomo, Development of a reporter peptide that catalytically produces a fluorescent signal through alpha-complementation. Protein Sci 24, 599–603 (2015).

15. D. M. Kar, I. Ray, in Data and Applications Security and Privacy XXXIII, S. N. Foley, Ed. (Springer International Publishing, Cham, 2019), pp. 61–80.

16. L. Adam et al., Strengths and limitations of the federal guidance on synthetic DNA. Nat. Biotechnol. 29, 208–210 (2011).

17. S. Galanie, K. Thodey, I. J. Trenchard, M. Filsinger Interrante, C. D. Smolke, Complete biosynthesis of opioids in yeast. Science 349, 1095–1100 (2015).

18. G. M. Church, Y. Gao, S. Kosuri, Next-generation digital information storage in DNA. Science 337, 1628 (2012).

19. Y. Erlich, D. Zielinski, DNA Fountain enables a robust and efficient storage architecture. Science 355, 950–954 (2017).

